# Pathotypr: harmonised MTBC lineage assignment and resistance-associated variant detection for genomic surveillance

**DOI:** 10.64898/2026.03.24.714002

**Authors:** Paula Ruiz-Rodriguez, Mireia Coscollá

## Abstract

**BACKGROUND:** Rapid, interoperable whole-genome tools for Mycobacterium tuberculosis complex (MTBC) surveillance remain limited for harmonised lineage assignment across recognised lineages and simultaneous resistance-associated variant detection.

**AIM:** To develop and validate Pathotypr, an alignment-free tool for harmonised MTBC lineage assignment and resistance genotyping from assemblies and raw reads.

**METHODS:** We reconstructed an MTBC phylogeny from 26,813 genomes using 609,003 polymorphic sites, derived an updated lineage marker backbone, and implemented a k-mer/Random Forest framework with marker-based lineage and WHO catalogue-based resistance calling. Performance was evaluated on 498 RefSeq assemblies, 88,071 UShER-TB typed sequencing samples, 162 clinical read sets for closest-reference matching, and 7,148 CRyPTIC isolates with phenotypic drug susceptibility data.

**RESULTS:** Pathotypr supported all 14 currently recognised MTBC lineages (L1-L10, A1-A4). On 498 complete genomes, marker-based and alignment-free lineage calls were 100% concordant, and prediction accuracy remained 100% on 254 independent assemblies. In 88,071 non-ambiguous UShER-TB samples, root-lineage concordance with TB-Profiler was 100%, while Pathotypr additionally identified lineage 10, A1 and A2. Resistance predictions showed 85.0% genotype-phenotype concordance overall, with high performance for rifampicin (95.8% sensitivity, 95.0% specificity) and isoniazid (93.0%, 97.9%). Runtime was about 1 second per sample, enabling analysis of 88,071 samples in approximately 24 hours on four threads. In the MDR-enriched CRyPTIC collection, Pathotypr supported reconstruction of 135 probable introduction events into Germany, Italy and Ukraine; 33.7% of introduction-associated isolates carried MDR/pre-XDR genotypes.

**CONCLUSION:** Pathotypr enables rapid, harmonised MTBC lineage assignment and high-confidence resistance screening, supporting near real-time and cross-border tuberculosis surveillance.

## Introduction

*Mycobacterium tuberculosis* remains the leading cause of death from a single infectious agent, causing more than one million deaths each year [1]. In tuberculosis control, sequencing is useful only if results are returned rapidly enough to inform outbreak detection, contact tracing, transmission monitoring, and timely identification of resistance-associated variants. Yet genome-based tuberculosis surveillance still covers only a small proportion of cases, despite the rapid adoption of near real-time sequencing for SARS-CoV-2.

This gap is increasingly problematic because MTBC classification is shifting from species-based nomenclature to lineage-based genomic classification. The *Mycobacterium tuberculosis* complex (MTBC), once regarded as genetically monomorphic, is now recognised to comprise ten human-adapted lineages (L1-L10) and four animal-adapted clades (A1-A4) [2–6]. Genomic collections remain uneven, with strong geographic and host bias. African lineages, especially L5-L10, and MTBC from animal reservoirs are under-represented, reducing the ability to detect uncommon introductions and complicating interpretation of zoonotic and anthropozoonotic transmission.

MTBC lineage assignment has shifted from targeted genotyping to whole-genome sequencing, with SNP phylogenies now serving as the reference standard. Routine workflows rely on SNP-based callers with complementary strategies: KvarQ screens raw FASTQ reads for predefined markers using curated test suites [7], whereas SNP-IT and TB-Profiler typically map reads to a reference genome and interrogate diagnostic SNPs to assign subspecies and lineages while also reporting resistance-associated mutations [8–10]. In practice, lineage assignment and resistance profiling are often interpreted together, particularly when investigating the emergence or spread of drug-resistant clusters. However, four issues continue to limit comparability and complicate surveillance interpretation.

First, inconsistent naming across studies and databases makes MTBC surveillance data difficult to compare. Lineages 5 and 6 are frequently reported as *M. africanum*, while animal-adapted clades are often treated as separate species: A1 (chimpanzee bacillus, dassie bacillus, *M. mungi*, *M. suricattae*), A2 (*M. microti*, *M. pinnipedii*), A3 (*M. orygis*), and A4 (*M. bovis*, *M. caprae*) [2]. Zwyer et al. [3] proposed a livestock-associated scheme (La1-La3) that groups animal-adapted diversity and excludes A1, further complicating comparisons across laboratories and over time. Second, barcode-based typing depends on timely updates as the MTBC phylogeny expands. Newly described lineages such as L10 are not consistently represented in reference tools such as TB-Profiler because an official marker set has not yet been released [4]. Third, misclassification is documented: Ngabonziza et al. [6] described lineage 8 and reported one sequence misclassified as *M. bovis* in public draft genome collections. Fourth, current tools do not robustly support lineage typing from complete genome assemblies that differ in coordinate systems or reference orientation.

To address these limitations, we developed Pathotypr, an open-source tool for harmonised MTBC lineage assignment and resistance-associated variant detection from reads and assemblies. Pathotypr implements an updated SNP scheme covering all recognised lineages and combines alignment-free k-mer counts with a Random Forest model to avoid reference-mapping bottlenecks. It can also type genomes lacking canonical markers. Here, we describe Pathotypr and evaluate its performance across heterogeneous genomic inputs relevant to routine surveillance. Designed for use on standard desktop hardware (<8 GB RAM) and available through both command-line and graphical interfaces, Pathotypr enables interoperable, near real-time reporting across laboratories and supports cross-border public health surveillance.

## Methods

### Study design and datasets

We developed and validated Pathotypr as an alignment-free framework for harmonised *Mycobacterium tuberculosis* complex (MTBC) lineage assignment, detection of resistance-associated variants, and closest-reference selection from genome assemblies and raw sequencing reads. The study comprised four stages: reconstruction of an updated MTBC phylogenetic backbone, derivation of lineage-defining SNP markers, implementation of marker-based and k-mer-based classification modules, and evaluation on independent validation, benchmarking, closest-reference, and clinical datasets. Pathotypr was designed to resolve all currently recognised human-adapted lineages (L1-L10) and animal-adapted clades (A1-A4). The datasets analysed included 26,813 genomes for phylogenetic reconstruction (available in Supplementary Table 1), 498 complete RefSeq genomes for lineage validation, 254 chromosome-level assemblies withheld from model training for independent testing (Supplementary Table 2), 88,071 UShER-TB samples after exclusion of mixed infections (Supplementary Table 3), 162 paired-end read sets for closest-reference evaluation (Supplementary Table 4), and 7,148 CRyPTIC isolates with linked phenotypic drug susceptibility testing (pDST) data and epidemiological metadata (Supplementary Table 5).

Genome sources, accession lists, and dataset-specific inclusion and exclusion criteria are provided in aforementioned Supplementary Tables.

### Phylogenetic reconstruction and lineage marker derivation

Polymorphic sites were extracted relative to the MTB_ancestor reference using SNPick v0.1.0 (https://github.com/PathoGenOmics-Lab/snpick). Sites previously reported as problematic by Modlin et al. [11], together with positions listed in the WHO mutation catalogue as associated with drug resistance, were removed before phylogenetic inference to minimise artefacts caused by recurrent mutation. The final alignment comprised 609,003 polymorphic sites across 26,813 genomes. Maximum-likelihood phylogenetic inference was performed with IQ-TREE v2.0.0 [12] under the GTR+G4 model with ascertainment bias correction. Phylogenies were visualised and annotated using custom Python scripts and coloured with the mycolorsTB palette (https://github.com/PathoGenOmics-Lab/mycolorsTB).

Branch-specific substitutions were reconstructed with TreeTime v0.11.4 [13]. Candidate lineage markers were defined as SNPs located on the stem branch leading to each monophyletic lineage or clade. Variants detected both on a lineage-defining branch and elsewhere in the tree were classified as homoplastic and excluded. Remaining variants were functionally annotated against the H37Rv reference genome (GenBank accession AL123456.3) using SnpEff v5.1 [14], and only non-missense, non-homoplastic variants were retained in the final marker panel. The final Pathotypr marker set comprised 2,145 lineage-defining SNPs. Lineage reporting followed a harmonised nomenclature in which human-adapted groups were designated L1-L10 and animal-adapted groups A1-A4.

### Pathotypr implementation and resistance genotyping

Pathotypr was implemented in Rust as both a standalone command-line tool and a cross-platform desktop application built with Tauri. The software comprises five modules: Train, Predict, Classify, Split-FASTQ, and Match.

The Train module converts labelled assemblies into feature-hashed sparse k-mer vectors projected into 2²⁰ hash buckets and fits a Random Forest classifier with 100 trees and unweighted Gini impurity using a single 80:20 train–test split (seed = 42). The final model used k = 31, selected on the basis of classification accuracy over k = 21. The Predict module applies the trained model to new assemblies for alignment-free lineage prediction.

The Classify module screens assembled genomes for lineage-defining SNP markers using diagnostic 31-bp k-mers and assigns the deepest lineage supported by a complete ancestral path from root to tip. Ties are resolved by SNP count and then lexicographic order. Profiles lacking a valid nested path are assigned to the most abundant lineage overall, while genomes without marker hits are reported as "Unclassified".

The Split-FASTQ module performs direct k-mer-based genotyping from raw FASTQ(.gz) reads. Variants at marker positions are called when read depth is at least 10× and alternate allele frequency is at least 95%. Resistance-associated variants were screened against the World Health Organization mutation catalogue, second edition (2023), restricted to confidence grades 1 and 2. Detected variants were reported with genomic position, nucleotide and amino acid change, associated drug, and WHO confidence grade. Per-drug genotypic calls were aggregated into standard resistance categories: pan-susceptible (Pan-S), isoniazid mono-resistant (INH-R), other resistance (Other-R), multidrug-resistant (MDR), and pre-extensively drug-resistant (pre-XDR).

The Match module compares sample k-mer profiles against a curated panel of representative reference genomes and returns the closest reference using a weighted k-mer containment score in a two-stage coarse/fine search with optional early stopping.

### Validation, external benchmarking, and closest-reference evaluation

Lineage assignment was first assessed on 498 complete RefSeq genomes by comparing Classify and Predict outputs with phylogenetic placement. Generalisability was then evaluated on 254 chromosome-level assemblies withheld from model training. External benchmarking used 88,071 UShER-TB samples [15] after exclusion of mixed infections. Concordance with TB-Profiler was assessed at the root-lineage level using predefined equivalence rules for animal-adapted nomenclature differences (La1 = A4, La2 = A2, La3 = A3). Groups absent from the comparator marker scheme, including L10, A1, and A2, were analysed separately.

Runtime and memory usage were measured on a Mac mini M2 (8 GB RAM) running macOS Sequoia with four threads. For closest-reference evaluation, 162 paired-end clinical read sets spanning 13 lineages were mapped both to MTB_ancestor and to the Pathotypr-selected closest reference using bwa-mem2 v2.3. Variants were called with bcftools mpileup and call v1.22. Mapping performance was summarised by breadth of coverage, per-base error rate, lineage-to-reference concordance, and the number of non-fixed SNPs, defined as positions with allele frequencies between 0.05 and 0.90.

### CRyPTIC application and statistical analysis

A subset of 7,148 *M. tuberculosis* isolates from the CRyPTIC consortium [16] with linked pDST data and epidemiological metadata, including country of isolation, subnational region, and collection year (2001-2019), was analysed using the Split-FASTQ workflow. Resistance mutations were called using the same thresholds described above and restricted to WHO confidence grades 1 and 2. Equivalent phenotypic resistance categories were derived from binary pDST results using the same hierarchical definitions. Per-drug performance was assessed by sensitivity, specificity, positive predictive value, and negative predictive value, with exact denominators reported. Concordance between genotypic and phenotypic resistance categories was calculated as the proportion of isolates assigned to the same ordinal category, and discordance was graded by ordinal distance (1, minor; 2, moderate; ≥3, severe).

A maximum-likelihood phylogeny was reconstructed from whole-genome SNP alignments using IQ-TREE v2.0.0 under the GTR+G4 model. The tree was rooted on the most recent common ancestor of eight L6/*M. africanum* and animal-lineage isolates, representing the deepest-branching clade within the MTBC. Pathotypr lineage assignments, genotypic resistance profiles, country of isolation, collection year, and phenotypic resistance category were projected onto the phylogeny for integrated visualisation.

Ancestral geographic states were inferred with TreeTime mugration using ISO 3166-1 alpha-3 country codes as a discrete trait. Probable independent introduction events into Europe were defined as phylogenetic transitions in which the inferred parent node was non-European and the descendant node was assigned to Germany, Italy, or Ukraine. For each introduction event, all descendant tips were collected and their genotypic resistance categories tallied. Clades containing more than 500 tips were excluded to reduce artefactual deep-branching assignments. Introduction routes were aggregated by donor-recipient country pairs. Enrichment of MDR/pre-XDR genotypes among introduced versus autochthonous European isolates was tested using two-sided Fisher’s exact test, with odds ratios and exact p-values reported. Statistical analyses, data integration, and figure generation were performed in Python v3.12 using matplotlib v3.10, scipy v1.15, and dendropy v5.0.

## Data availability

The Pathotypr source code and documentation are available at https://github.com/PathoGenOmics-Lab/pathotypr.

The phylogenetic marker sets used for lineage classification are deposited in Zenodo (https://doi.org/10.5281/zenodo.19210044). Sequencing data analysed in this study are publicly available from the NCBI Sequence Read Archive and RefSeq; accession numbers for all samples are listed in Supplementary Tables 1–5.

## Results

### An updated phylogenetic marker backbone

To establish the phylogenetic framework implemented in Pathotypr, we reconstructed a large-scale maximum-likelihood phylogeny from 26,813 *Mycobacterium tuberculosis* complex (MTBC) genomes using 609,003 polymorphic sites. This comprehensive dataset captured the currently recognised diversity of the MTBC and resolved the 10 human-adapted lineages (L1-L10) and four animal-adapted clades (A1-A4) (Figure 1A). Sampling was highly uneven across groups, reflecting the known structure of available genomic data. Most genomes belonged to L4 (n=11,237) and L2 (n=6,273), whereas recently described or geographically restricted lineages were represented by few sequences, including L8 and L10 (n=2 each), L9 (n=5), and L7 (n=29). Animal-adapted diversity was similarly imbalanced, with A4 comprising 4,412 genomes compared with 9 in A1, 28 in A2, and 55 in A3. Nevertheless, inclusion of all available groups enabled construction of a unified phylogenetic backbone spanning both common and rare MTBC diversity.

**Figure 1.**
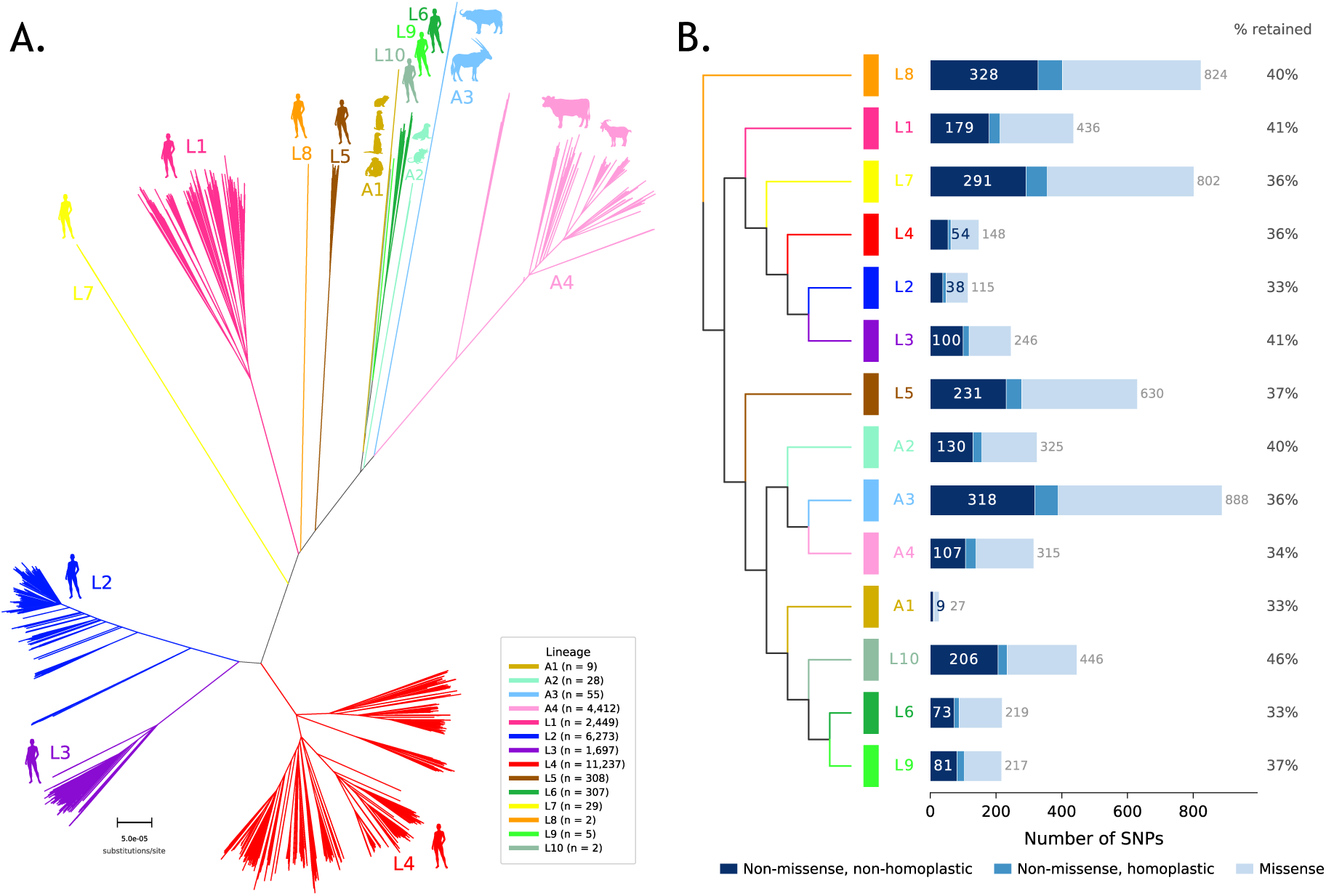
Phylogenetic framework of *M. tuberculosis* and selection of Pathotypr phylogenetic markers. **(A)** Maximum-likelihood phylogeny inferred from 26,813 genomes and 609,003 polymorphic sites. Branches are coloured by human-adapted lineages (L1-L10) and animal-adapted clades (A1-A4); typical hosts are indicated as silhouettes and sample sizes are shown in the legend. Scale bar, substitutions per site. **(B)** Lineage-collapsed cladogram summarising SNP selection on the stem branch leading to each lineage or clade. Stacked bars show candidate lineage-leading SNPs partitioned into missense variants, homoplastic non-missense variants, and retained non-missense, non-homoplastic markers; retained proportions are shown at right.

We then derived candidate phylogenetic markers for Pathotypr by mapping variants onto the phylogeny and extracting SNPs located on the stem branch leading to each monophyletic lineage or clade (Figure 1B). This phylogeny-informed strategy prioritised markers reflecting shared evolutionary ancestry rather than recent or recurrent variation. Across the phylogeny, we identified 5,638 candidate SNPs, of which 3,039 missense variants and 454 homoplastic non-missense variants were excluded, yielding 2,145 markers for lineage assignment in Pathotypr. Retention rates were similar across lineages and clades, ranging from 33% in Lineages 2, 6 and A1 to 46% in Lineage 10. These lineage-leading, non-missense, non-homoplastic SNPs define the updated phylogenetic marker set implemented in Pathotypr. However, due to the underrepresentation, performance for L5 to L10 and animal lineages should be interpreted cautiously.

### A new tool to predict, classify and match genomes

Pathotypr informs of i) lineage assignment and drug resistance signatures of genomic assemblies, ii) genotypes lineage defining and drug resistance markers for genome assemblies and short reads, and iii) return the closest assembled genome of short reads sequences (Figure 2A). The toolkit integrated five modules: 1. *Train*, which builds Random Forest classifiers from labelled genome assemblies; 2. *Predict*, which assigns lineages to new assemblies using the trained model; 3. *Classify*, which screens assemblies against a curated set of lineage-defining and drug-resistance SNP markers; 4. *Split-FASTQ*, which genotypes lineage and resistance directly from raw FASTQ reads without alignment; and 5. *Match*, which identifies the most similar reference genome for each query by streaming k-mer comparison. On desktop hardware, training on 10 complete genomes took 0.61 s with 302 MB RAM and scaled to 55 s and 1.4 GB for 50 genomes. Prediction classified five genomes in 0.25 s, screening 3,707 SNPs across five assemblies took 0.10 s, and direct genotyping of a 65× paired-end sample from FASTQ took 10.5 s with constant memory use of about 26 MB. In the CRyPTIC dataset, Pathotypr processed 7,148 samples at a mean of 4.97 s per sample and 196.7 MB RAM, corresponding to about 362 samples per hour on a standard workstation.

**Figure 2.**
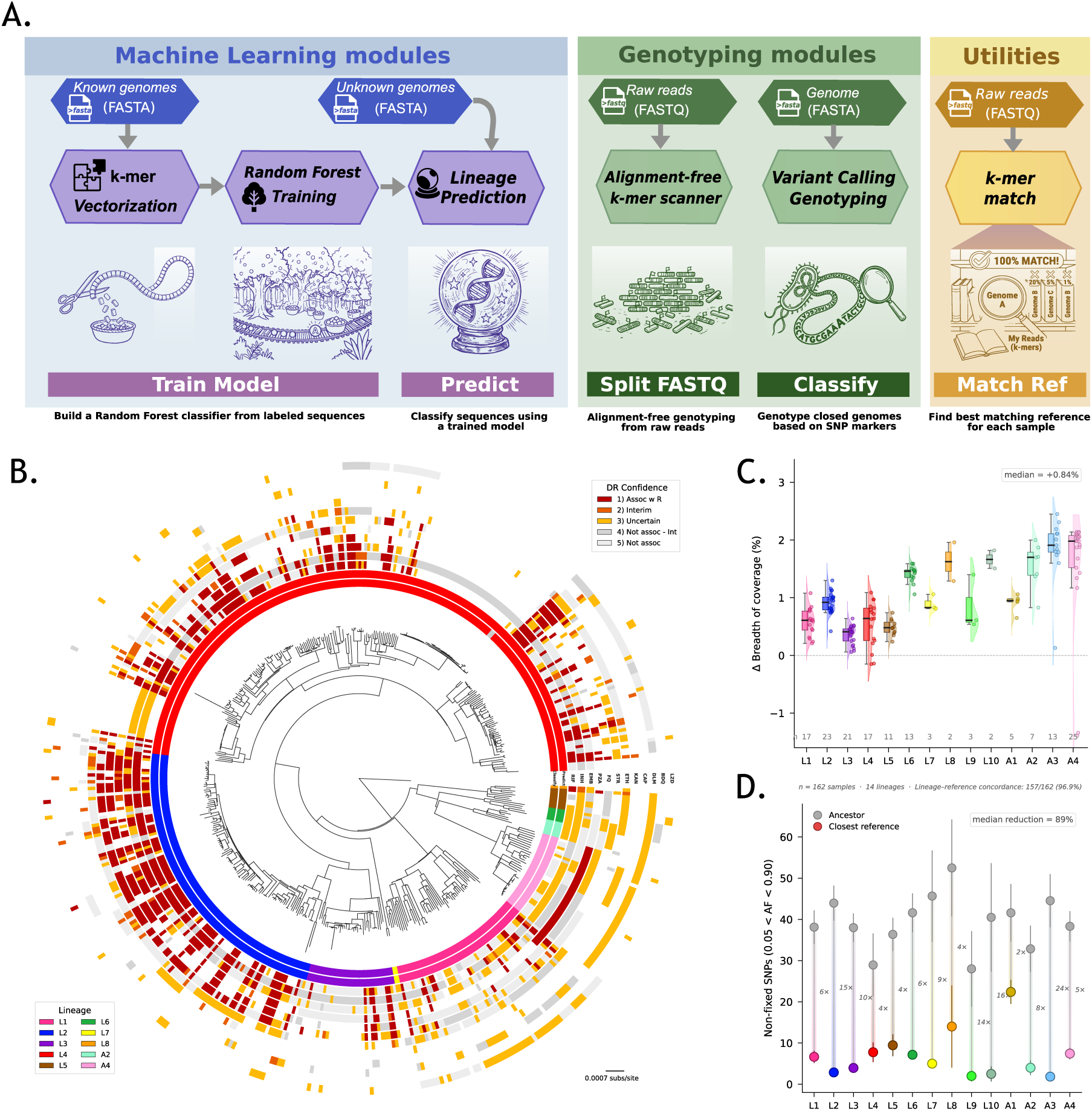
Overview and validation of Pathotypr for alignment-free lineage classification and genotyping of *M. tuberculosis* complex genomes. **(A)** Schematic overview of the five Pathotypr modules grouped into three categories: machine learning, genotyping and utilities. Train builds a random forest classifier from k-mer-vectorised labelled genome assemblies; Predict assigns lineages to unknown genomes; Split-FASTQ performs alignment-free genotyping from raw sequencing reads; Classify calls user-defined single nucleotide polymorphism (SNP) markers on assembled genomes; and Match Ref identifies the closest reference genome from a candidate panel by k-mer comparison. **(B)** Maximum-likelihood phylogeny of 498 *M. tuberculosis* complex genomes rooted on lineage 8. Scale bar, substitutions per site. The inner ring shows lineage assignments from Classify and the second ring shows predictions from Train/Predict (random forest, k = 31). Both methods showed 100% concordance across all 498 classifiable genomes. Outer rings show resistance profiles for 12 antimicrobials, coloured by World Health Organization confidence level for association with resistance. **(C)** Raincloud plots showing the improvement in breadth of coverage for 162 samples across 14 lineages when sequencing reads were mapped to the closest reference genome identified by Match Ref rather than to the *M. tuberculosis* ancestor. Mapping to the closest reference increased the breadth of coverage overall (median: +0.83 percentage points). **(D)** Dumbbell plot showing the reduction in non-fixed SNPs (0.05 < allele frequency < 0.90) when reads were mapped to the closest reference rather than to the ancestor. Numbers indicate the fold-reduction per lineage. Use of the closest reference reduced non-fixed SNPs by a median of 89%. Lineage-reference concordance was 95.6% (155/162); the five discordant samples belonged to lineage A1, for which no representative reference was available in the panel.

We evaluated lineage assignment on 498 complete MTBC assemblies from NCBI RefSeq (Figure 2B) with *Classify* and *Predict* modules. Both assigned major lineages to 497/498 genomes, and all classifiable genomes matched their position on the maximum-likelihood phylogeny and the alignment-free Predict module showed the same result at k = 31. When trained on 398 genomes in an 80:20 split with Train module, the Random Forest model achieved 100% concordance with marker-based calls across the same 498 genomes. Accuracy of the Train module fell from 100% to 96.4% when reducing kmer size from 31 to 21. On an independent set of 254 chromosome-level assemblies excluded from training, Predict maintained 100% accuracy at kmer size of 31.

Pathotypr screened drug-resistance mutations using the WHO catalogue of mutations associated with drug resistance in *M. tuberculosis*. Among the 498 complete genomes, 256 (51.2%) carried at least one resistance-associated mutation graded 1 or 2 by the WHO catalogue. This high proportion does not reflect global DR-TB prevalence (∼3.3% MDR among new cases; [1]) but rather the well-documented sequencing bias toward clinically relevant strains. A single BioProject (PRJNA555636), explicitly focused on MDR/XDR-TB, contributed 92 genomes (18.4%), of which 79 (86%) carried confirmed resistance mutations. Lineage 2 showed the highest resistance rate (76%), consistent with its known association with MDR/XDR outbreaks, while the 32 *M. bovis*/BCG genomes (lineage A4) carried the intrinsic *pncA* H57D mutation conferring pyrazinamide resistance (Figure 2B).

To evaluate the Match module, we predicted the closest assembly to mapped paired-end reads from 162 clinical samples spanning 14 lineages typed with Split-FASTQ and then compared mapping and calling statistics between the closest predicted assembly to the gold standard reference *M. tuberculosis* ancestor [17] (Figure 2D). Because nearest-reference selection improved interpretation of read-based variant calls in a strongly structured pathogen like *M. tuberculosis [18,19]*, we expect that mapping to the inferred closest reference will improve mapping and calling statistics. First we confirmed Lineage concordance between sample and closest assembly in 96.9% (155/162); the five discordant cases all belonged to A1, for which no representative assembly was available in the panel. Then, we corroborated that mapping and calling statistics improved using the closest reference. It provided increased coverage breadth by a median of 0.83 percentage points and reduced error rates in calling fixed and non-fixed SNPs across all lineages, with the largest gains in A3 (+1.85), A4 (+1.44) and L6 (+1.41). It also reduced the number of non-fixed SNPs (0.05 < allele frequency < 0.90) by a median of 89% across lineages, decreasing the typical count from 20-60 to 1-15 (Supplementary Table 4). These results indicate that much of the apparent within-host heterogeneity seen against a distant reference reflects fixed inter-lineage divergence. Drug resistance profiling of the same 162 samples with the Split-FASTQ module detected 151 WHO grade 1–2 mutations across 13 drug categories (Supplementary Table 6). When resistance calls were restricted to high-confidence mutations (WHO grades 1 and 2), 99 isolates (61.9%) were classified as pan-susceptible, 18 as MDR, 8 as INH-R, 6 as pre-XDR/XDR, 3 as RR, and 26 as Other-R. The predominance of pan-susceptible isolates is consistent with the broad lineage diversity of the panel, which was designed to maximise phylogenetic representation rather than enrich for resistant strains.

### Large-scale lineage classification benchmark and comparison with gold standard tools

Together with TB-profiler [10], KvarQ [7] and Mykrobe [20], Pathotypr provides a graphical interface making it easily accessible (Table 1). It can operate not only in FASTQ, like most of the available tools, but it can also use genome assemblies, like SNP-It [8] and fastlin [21]. Similarly to others, Pathotypr also incorporates drug resistance genotyping. However pathotypr is the only tool that incorporates all available MTBC Lineages (Table 1), and it can also be easily escalated to incorporate new markers even to be used for other pathogens. Finally, Pathotypr is the quickest tool available (Table 1). On four threads, pathotypr processed 88,071 clinical samples in approximately 24 hours (∼1 s per sample), compared with an estimated 636 days for TB-Profiler under equivalent conditions. These characteristics make pathotypr particularly suited to large-scale MTBC genomic surveillance, where comprehensive lineage resolution, resistance profiling and rapid turnaround are required simultaneously. To prove the large scale classification capabilities, we performed a two way comparison with TB-profiler, that also classifies root and sublineages, supports the majority of the lineages and predicts drug resistance.

**Table 1.** Comparison of selected tools for lineage classification and drug resistance prediction in the *M. tuberculosis* complex. Tools were compared based on published documentation and benchmark data. Pathotypr was the only tool supporting all 14 currently recognised MTBC lineages (L1-L10, A1-A4), including the recently described L10. Runtime estimates were derived from published benchmarks and documentation; Pathotypr was benchmarked on a Mac mini M2 using four threads. TB-Profiler and SNP-IT require alignment to a reference genome, whereas Pathotypr, fastlin and Mykrobe can analyse raw sequencing reads directly. DR: drug resistance; GUI: graphical user interface; MTBC: *Mycobacterium tuberculosis* complex; RF: random forest.

| Feature | pathotyper | TB-Profiler | Mykrobe | SNP-IT | fastlin | KvarQ |
| --- | --- | --- | --- | --- | --- | --- |
| **Approach** | Split k-mer + RF | SNP barcode | de Bruijn graph | SNP panel | Split k-mer | Targeted k-mer scanning |
| **Input formats** | FASTA, FASTQ(.gz) | FASTQ, BAM, VCF | FASTQ | FASTA, VCF | FASTA, FASTQ, BAM | FASTQ |
| **Alignment-free** | Yes | No | Yes | No | Yes | Yes |
| **Lineage resolution** | Root + sublineage | Root + sublineage | Root (limited sub.) | Root + sublineage | Root + sublineage | Root + sublineage |
| **Lineages supported** | L1–L10, A1–A4 | L1–L9, A4–A3 | L1–L6 | L1–L7, animal clades | L1–L9, A4 | L1–L7, A4 |
| **L10 support** | Yes | No | No | No | No | No |
| **Animal lineages** | A1–A4 | A3–A4 | Limited | All MTBC subspecies | A4 (*M. bovis*) | *M. bovis*, *M. caprae* |
| **DR prediction** | Yes | Yes | Yes | No | No | Yes |
| **Runtime per sample (a)** | ~1 s | ~3–10 min | ~3 min | ~1–2 min | <5 s | ~2 min (b) |
| **GUI** | Desktop app | Web | Desktop app | No | No | Desktop app |
| **Standalone binary** | Yes | No | No | No | Yes | No |
| **Language** | Rust | Python | Python/C++ | Python | Rust | Python/C |

We used the Split-FASTQ module in Pathotypr to classify and compare assignments of 88,071 *M. tuberculosis* complex (MTBC) sequencing samples from the UShER-TB dataset (Table 2). After excluding mixed infections, root-lineage assignments were completely concordant with TB-Profiler for all lineages covered by both methods (Table 2). Agreement was complete for the human-adapted lineages (L1-L9) and for the animal-adapted clades A3 and A4 included in TB-Profiler. Isolates assigned by Pathotypr to L10 (n=2), A1 (n=6) and A2 (n=23) were not directly comparable because these lineages are not represented in the TB-Profiler marker set. Of the two L10 isolates, one was classified by TB-Profiler as La1.5 (*M. bovis*), while the other remained unassigned. In these non-comparable groups, the mean number of lineage-defining markers detected per sample was 120 (range: 117-122) for L10, 143 (range: 1-312) for A1 and 119 (range: 3-187) for A2. Across the dataset, the mean number of lineage-defining markers detected per sample ranged from 63 for L9 to 289 for A3, in keeping with variation in marker density and sequence quality. Pathotypr processed samples at approximately 1 second per sample, enabling classification of the full dataset in around 24 hours on four threads.

**Table 2.** Concordance of lineage classification by Pathotypr and TB-Profiler in 88,071 *M. tuberculosis* complex sequencing samples. Samples from the UShER-TB dataset <u>(Karim et al. 2025)</u> with mixed infections were excluded. Concordance was assessed at the root-lineage level. Mean marker counts are shown as mean (range) lineage-defining SNPs per sample used in Pathotypr. Estimated runtimes were approximately 1 second per sample for Pathotypr and 10 minutes per sample for TB-Profiler using four threads. † Classified by TB-Profiler but not by Pathotypr. ‡ TB-Profiler lacks lineage 10, A1, A2 markers; one sample was classified as La1.5 (*M. bovis*) and one was unassigned. § Includes 51 *M. caprae* samples (TB-Profiler La2), counted as concordant within A4.

| Lineage | pathotyr (n) | TB-Profiler (n) | Concordance | Markers (mean, range) | pathotyr time | TB-Profiler time |
| --- | --- | --- | --- | --- | --- | --- |
| L1 | 9 012 | 9 012 | 100.0% | 172 (2–204) | 2.6 h | 64 d |
| L2 | 28 263 | 28 263 | 100.0% | 71 (1–94) | 8.3 h | 208 d |
| L3 | 8 967 | 8 967 | 100.0% | 73 (1–110) | 2.6 h | 64 d |
| L4 | 36 118 | 36 118 | 100.0% | 99 (1–162) | 10.4 h | 261 d |
| L5 | 370 | 370 | 100.0% | 238 (3–253) | 6 min | 3 d |
| L6 | 847 | 847 | 100.0% | 89 (7–111) | 15 min | 6 d |
| L7 | 39 | 39 | 100.0% | 273 (158–277) | 39 s | 6.5 h |
| L8 | 2 | 2 | 100.0% | 199 (143–255) | 2 s | 20 min |
| L9 | 24 | 24 | 100.0% | 63 (10–66) | 25 s | 4.2 h |
| **L10 ‡** | 2 | 0 | 0% | 120 (117–122) | 2 s | — |
| **A1 ‡** | 6 | 0 | 0% | 143 (1–312) | 6 s | — |
| **A2 ‡** | 23 | 0 | 0% | 119 (3–187) | 23 s | — |
| **A3** | 198 | 198 | 100.0% | 289 (46–311) | 4 min | 1 d |
| **A4 §** | 4 200 | 4 200 | 100.0% | 272 (24–299) | 1.2 h | 29 d |
| **Total** | **88 071** | **88 049** | **99.9%** |  | **~1 d** | **~636 d** |

### Pathotypr-defined lineages and resistance profiles identified MDR-enriched introductions into Europe

To illustrate the range of analyses that Pathotypr can support, we applied it to 7,148 isolates from the CRyPTIC collection and integrated lineage using TreeTime mugration. Although this dataset is enriched for drug-resistant isolates and is not intended to represent contemporary population-based sampling across countries, it provides a useful test case for assessing whether Pathotypr can recover biologically coherent lineage, resistance and geographic patterns within a single analytical framework.

Pathotypr generated complete lineage and genotypic resistance profiles for all 7,148 isolates with Split-FASTQ module. When mapped onto the maximum-likelihood phylogeny, lineage assignments were consistent with the underlying tree structure and clustered in monophyletic groups (Figure 3). The collection was dominated by lineage 4 (L4; 3,760/7,148, 52.6%) and lineage 2 (L2; 2,444/7,148, 34.2%), followed by L1 and L3 (468 each, 6.5%), with only a small number of isolates assigned to rarer groups such as A3, A4 and L6.

**Figure 3.**
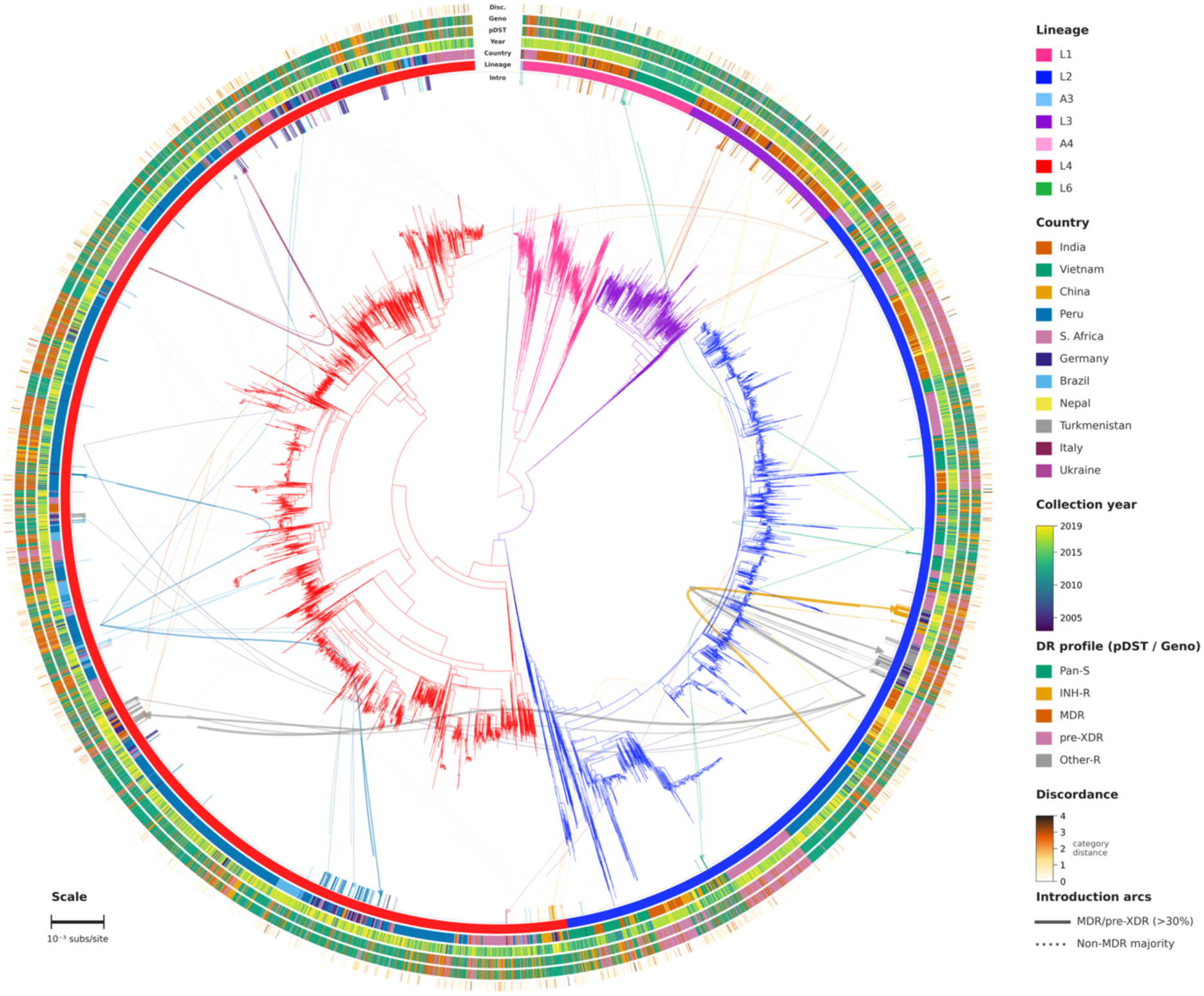
Circular phylogeny of 7,148 *M. tuberculosis* clinical isolates, CRyPTIC consortium, 2003-2019. Maximum-likelihood phylogeny reconstructed with IQ-TREE (GTR+G4), rooted using the lineage 6 (L6)/animal lineage outgroup and displayed in a circular layout. Branch colours indicate the major lineage assigned by Pathotypr. The six annotation rings show, from inner to outer, lineage, country of isolation, year of collection, phenotypic drug susceptibility testing profile, Pathotypr-predicted genotypic drug resistance profile based on World Health Organization (WHO) catalogue mutations (confidence grades 1 and 2), and discordance between phenotypic and genotypic resistance categories. Phenotypic profiles were classified as pan-susceptible (Pan-S), isoniazid mono-resistant (INH-R), multidrug-resistant (MDR), pre-extensively drug-resistant (pre-XDR), or other resistance (Other-R). Discordance is shown from white (distance = 0) to dark shading (distance = 4). Curved internal arcs indicate inferred cross-border introductions into Germany, Italy and Ukraine from ancestral-state reconstruction of country of origin using TreeTime mugration. Solid arcs indicate introduction events in which more than 30% of descendant isolates were MDR or pre-XDR; dotted arcs indicate introductions dominated by non-MDR isolates. Arc colour indicates the inferred source country. Tick marks at the periphery identify European isolates belonging to introduction clades and are coloured according to donor country. Scale bar, substitutions per site.

We then used the Pathotypr-annotated phylogeny together with ancestral-state reconstruction to identify 135 probable independent introductions from non-European source countries into Germany, Italy and Ukraine, involving 590 descendant isolates (Figure 4A). Most inferred introductions were into Germany, although this likely reflects the composition of the available dataset as well as the inferred phylogeographic structure. Introduction-associated clades were generally small: 69.6% were singletons and the median clade size was one isolate. Within the limits of this dataset, this pattern is more consistent with repeated detection of distinct imported strains than with large, sustained transmission chains following arrival, although the sampling framework does not allow direct inference

**Figure 4.**
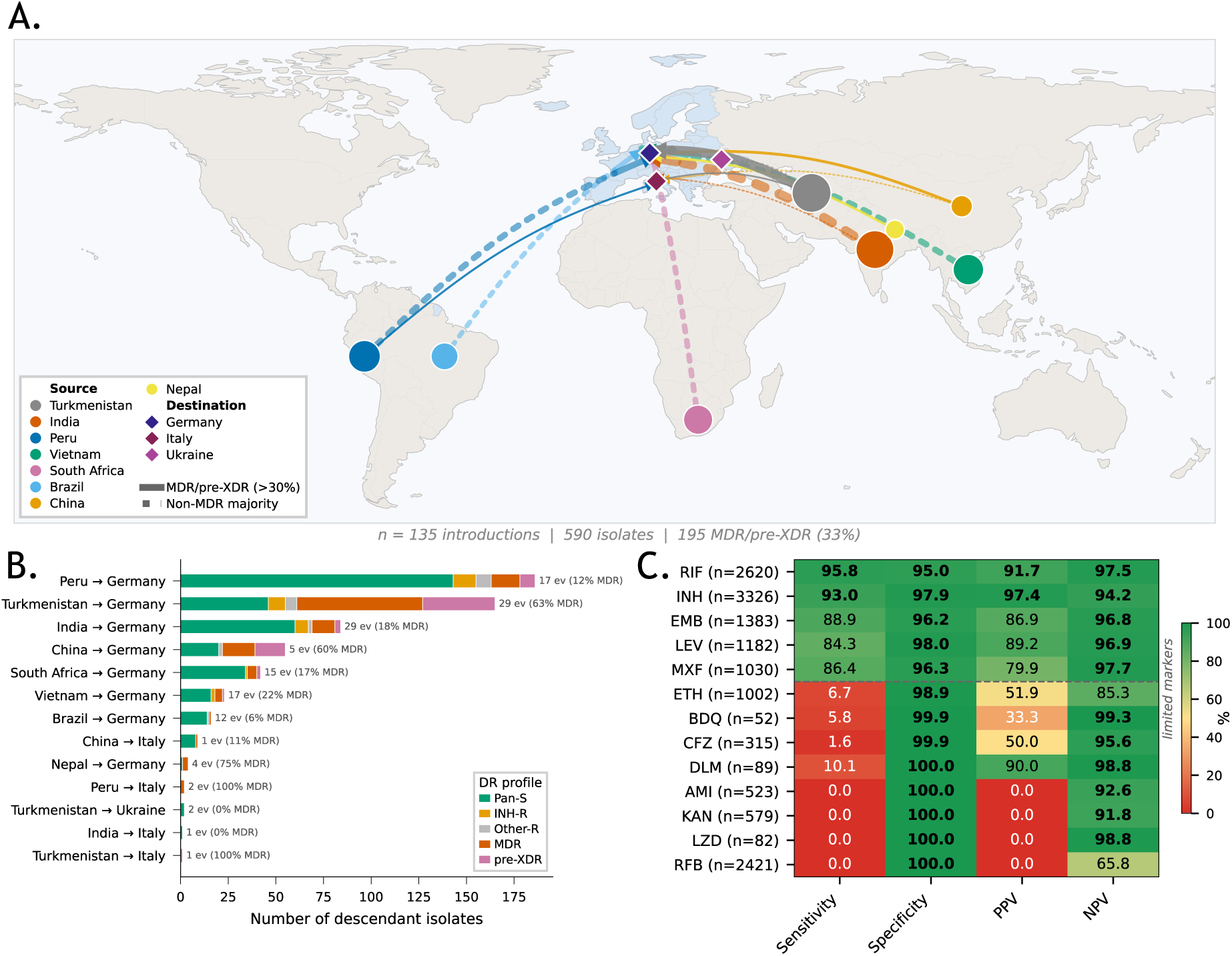
Cross-border introduction routes of *M. tuberculosis* into Europe and per-drug performance of Pathotypr genotypic resistance prediction. **(A)** World map showing inferred cross-border introduction events into Germany, Italy and Ukraine, based on ancestral-state reconstruction of geographic origin using TreeTime mugration across 7,148 fully annotated CRyPTIC isolates. Curved arrows connect source countries to European recipient countries. Solid arrows indicate routes in which more than 30% of descendant isolates were multidrug-resistant (MDR) or pre-extensively drug-resistant (pre-XDR), whereas dotted arrows indicate routes dominated by non-MDR isolates. Arrow width is proportional to the number of independent introduction events per route. Circles denote source countries and diamonds denote European recipient countries; Europe is highlighted in blue. Overall, 135 non-European introduction events comprising 590 descendant isolates were identified, of which 195 (33%) were MDR or pre-XDR. **(B)** Horizontal stacked bar chart showing the number of descendant isolates for each donor-to-recipient route, coloured by Pathotypr genotypic resistance profile: pan-susceptible (Pan-S), isoniazid mono-resistant (INH-R), other resistance (Other-R), MDR, or pre-XDR. Labels indicate the number of independent introduction events and the proportion of MDR/pre-XDR isolates per route. **(C)** Heatmap showing the performance of Pathotypr genotypic predictions based on World Health Organization (WHO) catalogue mutations (confidence grades 1 and 2) against CRyPTIC phenotypic drug susceptibility testing (pDST). Sensitivity, specificity, positive predictive value (PPV) and negative predictive value (NPV) are shown for 13 anti-tuberculosis drugs. The dashed line separates first-line drugs and fluoroquinolones from drugs with limited marker coverage in the current WHO resistance mutation catalogue. Drugs without Pathotypr markers, namely amikacin (AMI), kanamycin (KAN), linezolid (LZD) and rifabutin (RFB), showed 0% sensitivity, as expected.

about the relative contribution of importation and onward transmission at population level. The lineage composition of these inferred introduction events was also biologically plausible. Probable introductions from Turkmenistan and China were enriched for L2, whereas those from Peru and Vietnam were almost entirely L4; India contributed the greatest lineage diversity, including L1, L3 and L4 (Figures 3 and 4B). Because Pathotypr lineage calls remained congruent with the phylogeny, isolates assigned to the same broad lineage could still be resolved as separate introduction events when they occupied distinct branches. Within this illustrative analysis, introduced isolates carried a substantially higher resistance burden than other European isolates in the dataset. Overall, 33.7% of isolates assigned to probable introduction events had MDR or pre-XDR genotypes, compared with 5.5% of European isolates not assigned to external introductions. This enrichment was even more marked among L2 introductions, of which 73.4% were MDR or pre-XDR, compared with 20.5% of non-L2 introductions. At the dataset level, L2 also showed a higher phenotypic MDR rate than L4 (49.3% vs 28.4%; OR = 2.5; p = 5.36 × 10⁻⁶⁰; Fisher’s exact test). The highest MDR/pre-XDR proportions among introduction-associated clades were observed for Turkmenistan (62%) and China (53%) (Figure 4B). Because CRyPTIC is MDR-enriched, these findings should be interpreted as comparative patterns within the sampled collection rather than as estimates of the present-day burden of imported MDR-TB in Europe.

Across five resistance categories, genotypic and phenotypic classifications were concordant for 85.0% of isolates, and severe discordance was limited to 1.3%. Performance was highest for the two drugs defining MDR-TB, rifampicin (95.8% sensitivity, 95.0% specificity) and isoniazid (93.0%, 97.9%), and remained high for ethambutol (88.9%, 96.2%) and the fluoroquinolones levofloxacin (84.3%, 98.0%) and moxifloxacin (86.4%, 96.3%) (Figure 4C). Resistance was concentrated in a limited number of recurrent mutations: *rpoB* S450L accounted for 63.6% of rifampicin-resistant isolates, followed by D435V/Y (16.3%) and H445 variants (8.4%), while *katG* S315T was present in 76.6% of isoniazid-resistant isolates and inhA promoter variants in 29.6%. Among rifampicin-resistant isolates, 77.2% also carried *rpoC* or *rpoA* changes, consistent with compensatory evolution in transmissible MDR backgrounds. Fluoroquinolone resistance was mainly associated with gyrA substitutions at codons 90 and 94. By contrast, the absence of sensitivity for amikacin, kanamycin, linezolid and rifabutin reflected the lack of WHO grade 1 to 2 markers in the catalogue used here, while the lower sensitivity observed for ethionamide, bedaquiline, clofazimine and delamanid reflected the exclusion of lower-confidence variants (Supplementary Table 7, Figure 4C).

Taken together, these results show that Pathotypr can jointly resolve lineage, resistance and phylogenetic context within a single workflow, and that these annotations can be used to support cautious phylogeographic interpretation in complex tuberculosis datasets.

## Discussion

Pathotypr addresses a practical gap in tuberculosis genomic surveillance: the need for a fast, interoperable workflow that assigns *M. tuberculosis* complex (MTBC) lineages consistently across heterogeneous genomic inputs while also reporting resistance-associated variants. This is particularly relevant in Europe, where whole genome sequencing (WGS)-based TB surveillance is expanding but remains unevenly implemented. Surveys across EU/EEA countries have shown substantial heterogeneity in how molecular and genomic typing data are integrated into surveillance, and the ECDC pilot study demonstrated both the feasibility and the added value of standardised WGS-based surveillance for rifampicin-resistant and multidrug-resistant tuberculosis (MDR-TB) [22,23]. The 2025 ECDC molecular surveillance report further indicates that typing coverage remains suboptimal, despite continued detection of cross-border rifampicin-resistant and MDR-TB clusters [24]. Against this background, an updated Pathotypr backbone built from 26,813 genomes and spanning all currently recognised human-adapted and animal-adapted MTBC lineages represents a relevant contribution to harmonised cross-border reporting.

A major strength of Pathotypr is the combination of breadth, comparability and speed. Marker-based and alignment-free lineage assignments were fully concordant for all classifiable RefSeq assemblies, prediction accuracy remained 100% on independent held-out genomes, and root-lineage concordance with TB-Profiler was complete for all lineages represented in both tools in the 88,071-sample UShER-TB benchmark. The clearest added value was for groups that are incompletely represented in some existing tools, including Lineage 10 and the animal-adapted A1 and A2 clades. This is important because MTBC nomenclature continues to evolve, and surveillance workflows rapidly lose interoperability when marker schemes are not updated in parallel with phylogenetic knowledge. In this respect, Pathotypr offers more than a simple benchmark against an existing classifier. It provides a harmonised classification layer that can be applied across both common lineages and rarer clades within a single framework, while retaining a runtime of around 1 second per sample and avoiding the computational overhead of alignment-based preprocessing.

The resistance results are also encouraging from a surveillance perspective. Pathotypr performed best for rifampicin and isoniazid, the two drugs that define MDR-TB, and also showed strong performance for ethambutol and the fluoroquinolones. This pattern is consistent with the current state of MTBC resistance prediction, where genomic inference is most reliable when resistance is concentrated in a limited number of recurrent, high-confidence variants captured by curated catalogues [16,25,26]. In the CRyPTIC dataset, rifampicin and isoniazid resistance were dominated by canonical mutations in *rpoB*, *katG* and the *inhA* promoter, which helps explain why a marker-based approach achieved high sensitivity for the phenotypes most relevant to surveillance. The frequent co-occurrence of *rpoC* or *rpoA* changes among rifampicin-resistant isolates further supports the biological coherence of these resistant backgrounds, in line with evidence that compensatory evolution can enhance the fitness and onward transmission of resistant MTBC strains [27]. By contrast, the weaker performance observed for amikacin, kanamycin, linezolid, rifabutin and several newer or repurposed drugs should be interpreted mainly as a consequence of the intentionally conservative marker set used here, restricted to WHO confidence grades 1 and 2, rather than as a limitation of k-mer-based detection itself.

An additional practical contribution of Pathotypr is the nearest-reference module. Mapping reads to the closest reference increased coverage breadth and reduced apparent non-fixed variation by a median of 89%, particularly in phylogenetically distant clades. In MTBC, where deep lineage structure and reference-related blind spots can distort variant interpretation, this has direct implications for the analysis of within-host diversity, mixed infection and transmission clustering [11,19] . Rather than treating reference choice as a purely downstream technical step, Pathotypr incorporates it directly into the workflow. This is useful because it reduces the risk that fixed inter-lineage divergence is misinterpreted as biologically meaningful heterogeneity. The fact that all discordant lineage-to-closest assembly assignments were confined to A1, for which no representative reference was available, also underlines the need to expand reference panels as rarer MTBC groups become better represented.

The CRyPTIC application illustrates why these combined outputs matter. By placing 7,148 isolates within a single lineage, resistance and phylogenetic framework, Pathotypr supported an illustrative reconstruction of probable introduction events into Germany, Italy and Ukraine and showed that introduction-associated isolates in this dataset were enriched for MDR or pre-XDR genotypes. At the same time, most introduction-associated clades were singletons, a pattern that in this collection is compatible with repeated detection of distinct imported strains rather than extensive secondary transmission after arrival. However, because CRyPTIC is an MDR-enriched, non-population-based collection and includes only three European recipient countries, these findings should not be interpreted as direct estimates of current importation burden or transmission dynamics in Europe. Their value lies instead in showing that Pathotypr can generate the harmonised genomic outputs needed to support cautious phylogeographic interpretation and prioritisation of clusters for follow-up. This is relevant for European surveillance, where WGS is increasingly used to distinguish recent local transmission from new importations and to investigate international clusters [23,28,29].

This study has several limitations. Rare human-adapted lineages and animal-adapted clades remain under-represented in the phylogenetic backbone, so performance estimates for some groups should still be interpreted cautiously. Concordance with TB-Profiler was assessed at the root-lineage level, and equivalence at finer sublineage resolution should not be assumed. Resistance calling was intentionally conservative because only WHO confidence grades 1 and 2 were included, favouring specificity over sensitivity for several drugs. The CRyPTIC analysis was also limited by its non-population-based design and restricted geographic representation within Europe. Nevertheless, the overall picture is clear: Pathotypr combines broad MTBC lineage coverage, strong performance for the drugs most relevant to MDR-TB surveillance, low computational requirements, and deployment both as a command-line binary and a desktop GUI. Continued updating of marker sets, reference panels and resistance catalogues will be essential, but the current results suggest that Pathotypr is already well positioned for routine and cross-border TB genomic surveillance.

## Conclusion

Pathotypr is a rapid and accessible tool for harmonised MTBC lineage assignment and high-confidence resistance genotyping from assemblies and raw reads. Its updated phylogenetic framework covers all currently recognised MTBC lineages, showed complete concordance with comparator calls for all shared lineages in large-scale benchmarking, and extended classification to recently described or previously unsupported groups. In an illustrative application to the MDR-enriched CRyPTIC dataset, Pathotypr combined lineage, resistance and phylogeographic information within a single workflow, enabling cautious reconstruction of probable introduction events into three European countries and identifying enrichment of MDR/pre-XDR genotypes among introduction-associated isolates within this collection. Together with its availability as both a command-line tool and desktop GUI, these features support its use in routine and cross-border TB genomic surveillance.

## Acknowledgements

Computational analyses were performed on the Garnatxa high-performance computing cluster at the Institute for Integrative Systems Biology (I2SysBio), a joint research centre of the University of Valencia and the Spanish National Research Council (CSIC).

## Funding statement

This work was supported by grants PID2021-123443OB-I00 (MCIN/AEI/10.13039/501100011033; FEDER, EU "A way to make Europe") and PID2024-158536NB-I00 (MICIU/AEI/10.13039/501100011033; FEDER, EU "A way to make Europe").

## Conflict of interest

None declared.

## Authors’ contributions

PR-R conceived and developed the software, designed the methodology, performed all computational analyses and benchmarking, curated the datasets, created the figures, and wrote the manuscript. MC contributed to the conceptualization and methodology, supervised the project, provided resources and funding, and wrote the manuscript. Both authors read and approved the final version.

## Notes

### Competing Interest Statement

The authors have declared no competing interest.

