## Supplementary material for "Pathotypr: harmonised MTBC lineage assignment and resistance-associated variant detection for genomic surveillance": Caption Tables

### Supplemental Tables

**Supplementary Table 1. Metadata for 26,813 Mycobacterium tuberculosis complex isolates used for pathotypr development and validation.** The table lists 26,813 MTBC whole-genome sequencing samples encompassing all 14 currently recognised lineages. For each sample, the table reports the lineage assigned by pathotypr, the ENA/SRA run accession, isolation date (where available), country of isolation, country of patient birth (where available), and BioProject or BioSample accession. Lineages are colour-coded according to the mycolorsTB palette used throughout this study.

**Supplementary Table 2. Assemblies of *M. tuberculosis* complex genomes included in this study.** A total of 754 genomes were retrieved from NCBI RefSeq and grouped into two datasets: Complete (n = 498), comprising complete genome assemblies, and Chromosome (n = 256), comprising chromosome-level assemblies used as an independent validation set. For each genome, the table reports the RefSeq and GenBank accession numbers, organism name, strain designation, assembly name, assembly level, BioProject and BioSample identifiers, genome size (bp), GC content (%), number of scaffolds, and contig N50 (bp). Lineages were assigned using two complementary approaches implemented in pathotypr: SNP-based marker genotyping (classify module) and k-mer-based random forest prediction (predict module). Prediction confidence is reported as the proportion of trees in the random forest ensemble supporting the assigned lineage.

**Supplementary Table 3. Large-scale lineage concordance between pathotypr and TB-Profiler across 88,071 *M. tuberculosis* complex isolates.** Lineage assignments were compared for 88,071 MTBC isolates from the UShER-TB dataset (Karim et al., 2025). For each isolate, the table reports the ENA/SRA run accession, the root lineage and deepest sublineage assigned by pathotypr split-FASTQ, together with the corresponding SNP marker counts, and the lineage reported by TB-Profiler v2 in the UShER-TB supplementary data. Concordance was assessed at the root-lineage level after harmonising nomenclature (TB-Profiler La1 = A4, La2 = A2, and La3 = A3). In addition, 51 M. caprae isolates assigned to A4 by pathotypr and to La2 by TB-Profiler were considered concordant within the animal-adapted clade. Among isolates with a callable comparison, 88,039 were concordant; the remaining 31 belonged to lineages L10 (n = 2), A1 (n = 6), and A2 (n = 23), which are not supported by TB-Profiler and were therefore reported without a concordance call. Metadata columns include organism, country and continent of origin, sequencing platform, library layout, TB-Profiler drug-resistance classification, and median sequencing coverage, as reported in the UShER-TB dataset. Samples with mixed infections, defined as samples with two or more lineage calls in TB-Profiler, and samples unclassified by pathotypr were excluded.

**Supplementary Table 4. Benchmark comparing alignment to the closest reference and the ancestral reference in 162 samples.** Reads were first classified with pathotypr classify and then matched, using pathotypr match, to the most similar genome among 500 complete RefSeq assemblies. Reads were subsequently aligned with bwa-mem2 to both the selected closest reference and the ancestral M. tuberculosis reference (Comas et al., 2010; Zenodo 3497110). The table reports mapping metrics for each reference, including mapped reads (%), breadth of coverage, mean depth, and error rate, together with variant-calling metrics, including total SNPs, fixed SNPs (allele frequency, AF, ≥ 0.90), and non-fixed SNPs (AF 0.05-0.90). Delta values are reported as Δ = closest reference - ancestral reference. Positive Δ values for mapped reads and breadth indicate improved alignment to the closest reference, whereas negative Δ values for error rate and SNP counts indicate reduced noise.

**Supplementary Table 5. CRyPTIC dataset: phenotypic and genotypic drug-resistance profiles of 7,148 *Mycobacterium tuberculosis* clinical isolates.** This table summarises 7,148 isolates from the

Comprehensive Resistance Prediction for Tuberculosis: an International Consortium (CRyPTIC) project, collected between 2001 and 2019 across 13 countries. For each isolate, the table reports the sample identifier, ENA run accession, country, region, collection date, site, and library layout. Binary phenotypic drug-susceptibility results (S, susceptible; R, resistant; I, intermediate) and minimum inhibitory concentrations (MICs) are provided for 13 drugs: rifampicin, isoniazid, ethambutol, ethionamide, amikacin, bedaquiline, clofazimine, delamanid, kanamycin, levofloxacin, linezolid, moxifloxacin, and rifabutin. Lineages were assigned using pathotypr split-FASTQ based on SNP marker counts. Genotypic and phenotypic drug-resistance categories (Pan-S, INH-R, Other-R, MDR, pre-XDR) were compared, and each isolate was assigned a discordance level (Concordant, Minor, Moderate, or Severe) and a discordance distance. Drugs predicted to be resistant on the basis of WHO grade 1 or 2 mutations are listed for each isolate. For the 552 isolates involved in international transmission events, the table also includes the introduction event identifier, source and destination countries, clade size, and the fraction of MDR/pre-XDR isolates within each clade.

**Supplementary Table 6. Drug-resistance profiling of 162 *Mycobacterium tuberculosis* clinical isolates using pathotypr Split-FASTQ and WHO-graded mutations.** Drug resistance was assessed in 162 paired-end clinical read sets representing 13 MTBC lineages using pathotypr split-FASTQ and a curated marker panel derived from the WHO catalogue of mutations associated with drug resistance in M. tuberculosis. Resistance calls were restricted to mutations classified as WHO confidence grade 1 (associated with resistance) or grade 2 (associated with resistance, interim). Mutations classified as grades 3-5 were excluded from resistance classification but retained in the total mutation count. For each isolate, the table reports the sample identifier, ENA run accession, sublineage assigned by pathotypr, the closest reference genome selected by pathotypr match, and the study drug-resistance category: pan-susceptible, INH-R, RR, MDR, pre-XDR/XDR, or Other-R. Per-drug resistance status (R/S) is provided for 13 drug categories: rifampicin, isoniazid, ethambutol, pyrazinamide, streptomycin, fluoroquinolones, kanamycin, capreomycin, ethionamide, linezolid, bedaquiline/clofazimine, delamanid, and para-aminosalicylic acid. The second sheet lists the 151 individual grade 1-2 mutations detected, including the drug, gene, specific mutation, WHO grade, genomic position, reference and alternative alleles, read counts, and alternative allele fraction. Of the 162 isolates, 99 were classified as pan-susceptible, 26 as Other-R, 18 as MDR, 8 as INH-R, 6 as pre-XDR/XDR, and 3 as RR.

**Supplementary Table 7. Drug resistance-associated mutations detected by pathotypr in 7,148 *Mycobacterium tuberculosis* clinical isolates from the CRyPTIC dataset.** The table reports 6,094 unique mutations detected across 14 drug categories: rifampicin, isoniazid, ethambutol, ethionamide, fluoroquinolones, bedaquiline/clofazimine, delamanid, pyrazinamide, linezolid, streptomycin, kanamycin, capreomycin, para-aminosalicylic acid, and other targets. For each mutation, the table lists the associated drug, gene, amino acid or nucleotide change, WHO confidence grade (1-5), and WHO classification label. Detection frequency is reported as the number of isolates carrying the mutation (N detected), together with the number phenotypically resistant (N pheno R) and phenotypically susceptible (N pheno S) among isolates with available phenotypic data (N with phenotype). Positive predictive value (PPV, %) is calculated as the proportion of mutation-carrying isolates that were phenotypically resistant. Mutations were identified in 48 genes, most frequently *katG* (n = 331), *rpoB* (n = 320), *embC* (n = 313), *ethA* (n = 304), and *rpoC* (n = 292). WHO grade 1 and 2 mutations represent variants with the strongest evidence for resistance association, whereas grades 3-5 include variants with uncertain, insufficient, or no evidence of resistance association.
